# 40 Hz visual stimulation increases gamma-band functional connectivity beyond the visual cortex

**DOI:** 10.64898/2026.11.04.757576

**Authors:** Tianyu Ma, Qiang Sun, Tjaša Mlinarič, Marc M. Van Hulle

## Abstract

Visual stimulation at 40 Hz is being investigated as a non-invasive intervention for Alzheimer’s disease, but its network-level mechanisms in the human brain remain unclear. EEG functional connectivity measures interactions between brain regions and can be used to examine how sensory stimulation and cognitive engagement affect activity across regions. We recorded EEG from 29 healthy young adults during Regular 40 Hz flicker and Irregular 30–50 Hz flicker, each with and without a target-counting task (2 × 2 within-participant). Source-space connectivity between visual, parietal, and frontal regions was estimated with the debiased squared weighted phase-lag index (dWPLI). Regular stimulation increased 38–42 Hz connectivity for all three region pairs, with the largest increase between visual and parietal regions. The increase was specific to the stimulation frequency and was observed with two source reconstruction methods and in sensor space. The visual–parietal increase remained significant after correction for source leakage (spatial mixing between reconstructed sources) and after removal of the estimated stimulus-locked component, whereas the increases in visual–frontal and parietal–frontal connectivity did not. The concurrent task did not significantly modulate the stimulation effect for any region pair. These results show that 40 Hz visual stimulation increases gamma-band connectivity beyond the visual cortex. However, this increase may be mainly driven by the same external input affecting different regions. In addition, a simple cognitive task did not significantly affect the connectivity, suggesting that the connectivity change is to some extent robust to cognitive states.

## 1. Introduction

Modulating brain activity through rhythmic sensory stimulation is a promising area of research in non-invasive neuromodulation. Sensory stimulation at 40 Hz has attracted considerable attention because of its neuromodulatory effects in animal models of Alzheimer’s disease and it is now being tested as a non-invasive intervention in patients (Iaccarino et al., 2016; Adaikkan et al., 2019; Martorell et al., 2019; Chan et al., 2022; Murdock et al., 2024; Chan et al., 2025). In mouse models, the effects of 40 Hz stimulation were not limited to sensory cortex but also involved higher-order regions such as the hippocampus and prefrontal cortex (Adaikkan et al., 2019). Existing human studies have similarly shown that the neural effects of 40 Hz sensory stimulation can extend beyond early sensory regions. A long-term 40 Hz audiovisual stimulation study reported changes in resting-state fMRI functional connectivity related to visual network and between the visual network and other brain regions (Chan et al., 2022). Scalp and intracranial EEG recordings during acute visual stimulation showed 40 Hz responses at frontal and hippocampal sites, which the authors interpreted as propagation of the 40 Hz entrainment (Khachatryan et al., 2022). A recent intracranial EEG study further found that 40 Hz visual stimulation elicited 40 Hz responses in the hippocampus and temporal and frontal regions, while enhancing phase synchronization between the hippocampus and other cortical regions (Mlinarič et al., 2025).

However, a 40 Hz response at sites distant from early sensory regions does not by itself show that these regions interact. Power, signal-to-noise ratio (SNR), and inter-trial phase coherence (ITPC), the measures typically used to establish the 40 Hz response, are computed at each site separately and therefore cannot reflect whether regions interact. A response at the same frequency can be observed at distant sites because of volume conduction, or because each region responds to the same stimulus (Bastos and Schoffelen, 2016; Schoffelen and Gross, 2009). Functional connectivity analysis addresses this question more directly, because phase-based connectivity measures quantify whether the phase difference between regions is consistent across trials (Bastos and Schoffelen, 2016). During periodic stimulation, however, a consistent phase difference can also arise when two regions follow the same stimulus.

Two recent studies examined gamma-band connectivity during 40 Hz visual stimulation. An MEG study found that gamma connectivity between the visual cortex and parietal and frontal regions differed between 32, 40, and 48 Hz stimulation (Petro et al., 2024), and a source-space EEG study in older adults found broader and stronger gamma connectivity when the stimulation frequency matched the individual optimal frequency (Yoon et al., 2025). These studies compared stimulation frequencies, and it remains unclear how much of the connectivity during 40 Hz stimulation remains after correction for source leakage and removal of the stimulus-locked component. Periodic visual stimulation elicits steady-state visual evoked potentials (SSVEPs) in the visual cortex, while visual processing also involves parietal and frontal association regions (Norcia et al., 2015). Parietal regions are involved in visual selection and spatial attention, while frontal regions contribute to maintaining task goals and regulating attention (Corbetta and Shulman, 2002).

Interactions between these regions allow sensory information processing to adapt to behavioral goals and task demands (Vossel et al., 2014), and oscillatory synchronization between regions is associated with these processes (Siegel et al., 2008). Selective attention can also modulate the amplitude of steady-state visual evoked responses (Morgan et al., 1996), and a concurrent cognitive task has been shown to increase the spatial extent of the 40 Hz SNR (Khachatryan et al., 2022). If this increase reflects propagation, a concurrent task would also increase the stimulation effect on connectivity, particularly for connectivity with frontal regions. Alpha oscillations are associated with the selection of visual information and the suppression of irrelevant information, and alpha phase synchronization between visual, parietal, and frontal regions varies with visual attention (Jensen and Mazaheri, 2010; Lobier et al., 2018). Alpha-band connectivity can therefore provide an additional measure of task-related network modulation.

The present study tested these assumptions using EEG connectivity analysis. In a within-participant design with stimulation regularity (regular periodic stimulation and irregular control stimulation) and task condition (NoTask/TargetCounting) as factors, we measured phase-based connectivity between visual, parietal, and frontal regions in 29 healthy young adults. We tested three hypotheses. First, if regular 40 Hz stimulation spreads from the visual cortex to higher-order regions, it would increase 38–42 Hz connectivity between visual, parietal, and frontal regions, with effects remaining after correction for source leakage and removal of stimulus-locked activity (H1). Second, based on previous evidence that a concurrent cognitive task increases the spatial extent of the 40 Hz response (Khachatryan et al., 2022), we predict that TargetCounting would enhance the effect of regular stimulation on gamma-band connectivity relative to NoTask, particularly for connections involving frontal regions (H2). Third, given the established association between alpha-band synchronization and visual attention, we expected that TargetCounting would increase alpha-band connectivity (H3).

## 2. Materials and methods

### 2.1 Participants

Thirty healthy adults with normal or corrected-to-normal vision took part in the study. One participant was excluded from all analyses because of insufficient data quality. The remaining 29 participants were analyzed (15 female; range 21–34 years). All participants provided written informed consent before data collection. The study was approved by the Ethics Commission Research UZ/KU Leuven (approval no. S62547). All analyses included these 29 participants unless stated otherwise.

### 2.2 Visual stimulation and experimental conditions

Visual stimulation was delivered using a custom-built panel containing an 8 × 8 white LED matrix controlled by an Arduino Uno microcontroller (Fig. 1B). The matrix was embedded in a white enclosure to diffuse the light. A red LED at the center of the panel served as the fixation point. Participants sat approximately 55 cm from the device in a dimly lit room (Fig. 1A).

**Figure 1.**
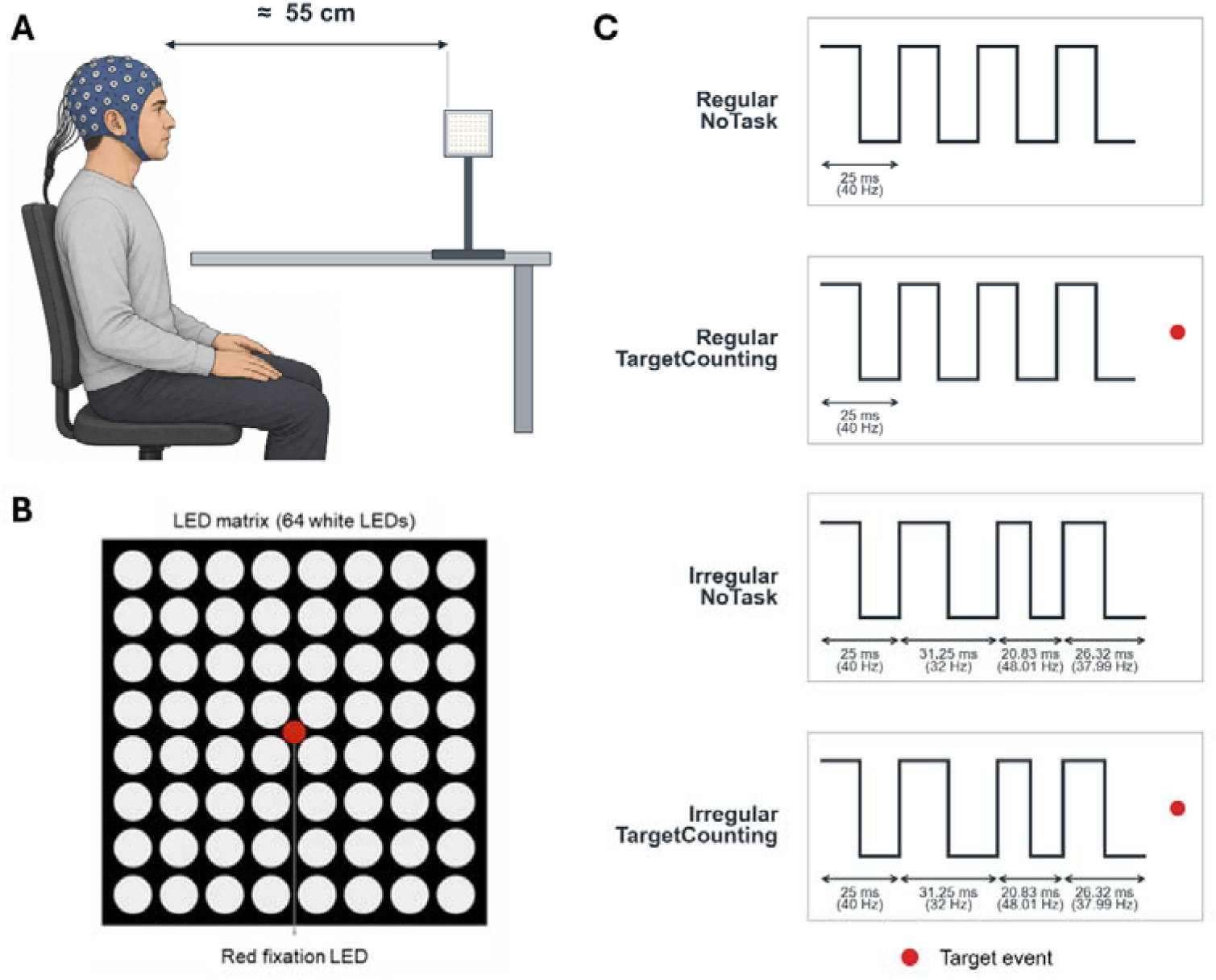
Experimental setup, visual stimulus, and experimental conditions. **(A)** Participants viewed the tabletop LED array from approximately 55 cm while EEG was recorded. **(B)** The device comprised an 8 × 8 array of 64 white LEDs and a central red fixation LED. **(C)** Four conditions crossed stimulation regularity (Regular or Irregular) with task condition (NoTask or TargetCounting). Regular stimulation used a constant 25-ms cycle (40 Hz). Irregular stimulation used variable cycle durations corresponding to instantaneous frequencies of 30–50 Hz; representative cycles are shown. In NoTask, participants viewed the array without an additional task. In TargetCounting, they counted brief illuminations of the red fixation LED (red dots). Panel **A** was generated using OpenAI’s image generation tool and edited by the authors.

The experiment used a 2 × 2 within-participant design. The two factors were stimulation regularity (Regular or Irregular) and task condition (NoTask or TargetCounting). Each participant completed four conditions: Regular–NoTask, Regular–TargetCounting, Irregular–NoTask, and Irregular–TargetCounting (Fig. 1C). Five minutes of stimulation were recorded per condition. The stimulation was presented either continuously or in two 2.5-min periods separated by a 1-min break. Regular stimulation consisted of 40 Hz square-wave flicker with a 50% duty cycle. Irregular stimulation consisted of consecutive square-wave periods with integer frequencies randomly selected between 30 and 50 Hz. The duty cycle and LED intensity (225-250 lumens) were the same in both conditions. During TargetCounting, the central red LED was illuminated for 250 ms at pseudorandom intervals centered on 30 s. Participants maintained fixation and mentally counted these events. During NoTask, participants maintained fixation without counting.

### 2.3 EEG acquisition and preprocessing

EEG was recorded at 1,000 Hz using a SynAmps RT system (Compumedics) using 62 scalp electrodes placed according to the extended international 10–20 system. FCz served as the online reference and AFz as the ground.

EEG data were processed in MATLAB R2021b using EEGLAB (development version, February 2026; Delorme and Makeig, 2004). Bad channels were detected automatically as channels with a flat signal or an outlying RMS amplitude (robust z-score > 6), using the data from all four conditions, and were reconstructed by spherical interpolation (15 of 29 participants; mean 1.3 channels, range 0–6). The data were filtered between 1 and 80 Hz, notch-filtered at 49–51 Hz, and rereferenced to the common average. Extended Infomax ICA was run once per participant on data pooled across conditions and temporarily downsampled to 250 Hz. Artifact-related components were identified automatically and removed based on their scalp maps, spectral properties, and kurtosis.

Five minutes of EEG, corresponding to the stimulation period, were analyzed for each condition. The data were divided into non-overlapping 4-s epochs. Epochs with invalid numerical values, or outlying amplitude or high-frequency power (robust z-score > 8, computed within each participant across all epochs of the four conditions) were excluded; the number of epochs was not equalized across conditions. The retained epochs were downsampled to 250 Hz for all further analyses; the downsampling before ICA was used only to estimate the decomposition. On average, 99.5%, 99.0%, 97.8%, and 96.4% of epochs were retained in the Regular–NoTask, Regular–TargetCounting, Irregular–NoTask, and Irregular–TargetCounting conditions, respectively (Friedman test, p = 0.050; Supplementary Table S1).

### 2.4 Source reconstruction and ROI time series

#### 2.4.1 Source reconstruction

Source reconstruction was performed using MNE-Python 1.12.1 (Gramfort et al., 2013) and the FreeSurfer fsaverage template (Fischl, 2012) provided with MNE-Python. Individual MRIs were not used; the template was used for all participants. The forward model used a three-layer boundary-element model and an ico-3 cortical source space. Source activity was estimated with a depth-weighted minimum-norm estimate (wMNE). The depth-weighting parameter was 0.8, the loose-orientation parameter was 0.2, and regularization was set to λ² = 1/9. Only the component normal to the cortical surface was retained.

#### 2.4.2 Regions of interest

Cortical regions were defined using the Desikan–Killiany atlas (Desikan et al., 2006). Regions of interest were selected a priori based on previous studies of 40 Hz visual stimulation (Khachatryan et al., 2022; Petro et al., 2024). The visual regions included the bilateral pericalcarine cortex, cuneus, and lingual gyrus. The parietal regions included the bilateral superior and inferior parietal cortex. The frontal regions included the bilateral caudal and rostral middle frontal cortex. Large regions were used because the spatial resolution of EEG source reconstruction is limited (Farahibozorg et al., 2018). Left- and right-hemisphere ROIs were analyzed separately (Fig. 2A).

**Figure 2.**
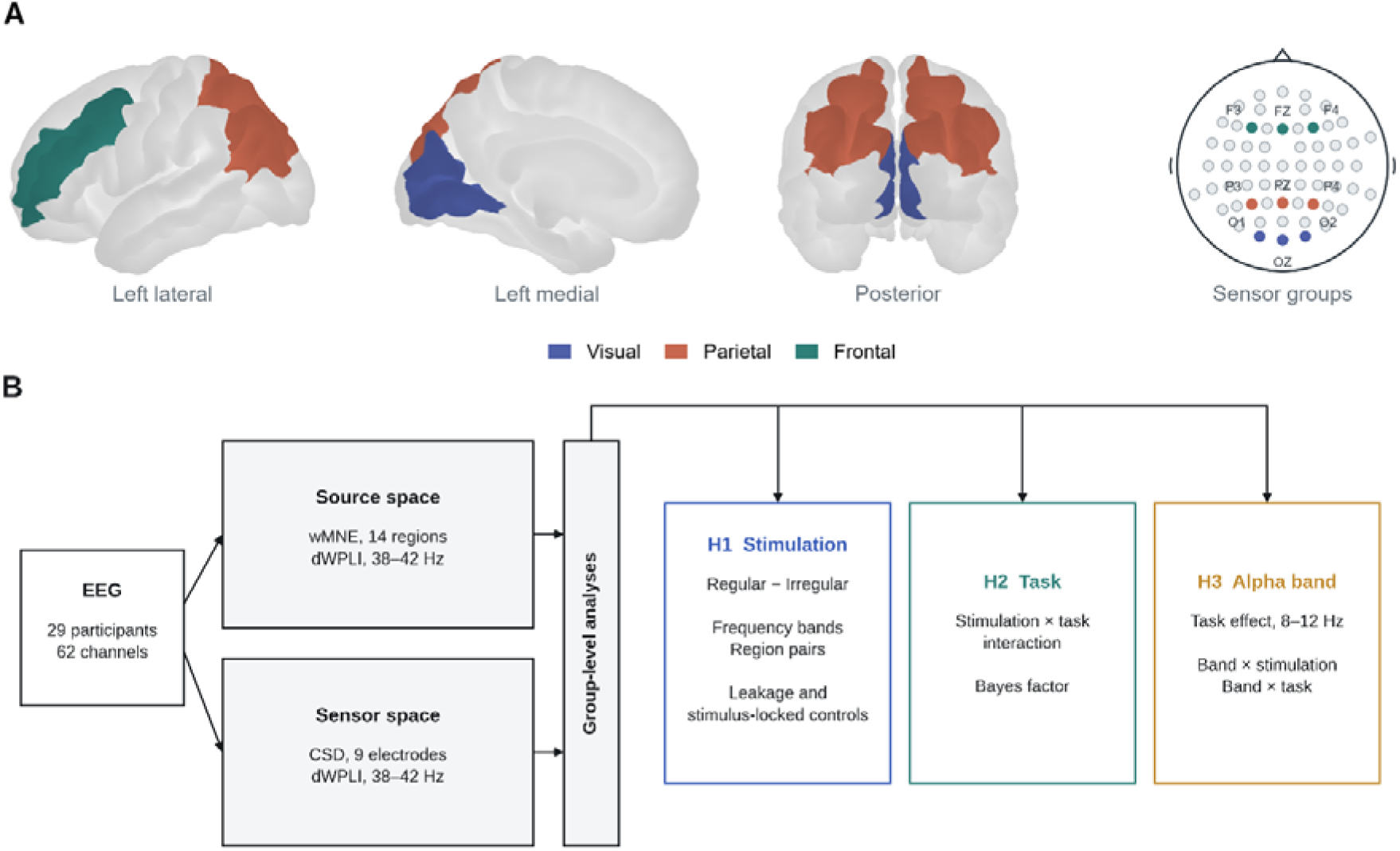
Regions of interest and analysis overview. **(A)** Visual, parietal, and frontal source-space regions and sensor groups. **(B**) Analysis overview: source- and sensor-space measures and the analyses used to test the three hypotheses (H1–H3). CSD refers to current source density.

#### 2.4.3 ROI time-series extraction

Source estimates were calculated for each 4-s epoch. One time series was extracted from each ROI using `pca_flip`, separately for each condition. This method uses the dominant principal component and aligns its sign according to the cortical surface orientation. The resulting ROI time series were used for the source-space connectivity analyses.

### 2.5 Connectivity estimation and experimental contrasts

#### 2.5.1 Connectivity estimation

The main analyses used the wMNE ROI time series in source space. Sensor-space analyses were used as sensitivity analyses. Source-space functional connectivity was calculated with MNE-Connectivity 0.8.1 (Fig. 2B) using the debiased squared weighted phase-lag index (dWPLI; Vinck et al., 2011). dWPLI reduces the contribution of phase differences near zero and corrects for bias related to the number of epochs. Cross-spectra were estimated from the 4-s epochs with a multitaper method (seven discrete prolate spheroidal sequence tapers, spectral smoothing ±1 Hz, frequency resolution 0.25 Hz; tapers weighted equally), and dWPLI was averaged across 38–42 Hz.

dWPLI was calculated for every pair of ROI time series in each condition and then averaged across all ROI pairs between visual and parietal (24 pairs), visual and frontal (24 pairs), and parietal and frontal regions (16 pairs).

#### 2.5.2 Experimental contrasts and comparisons

Let *C_RN_*, *C_RT_*, *C_IT_* and *C_IT_* denote connectivity in the Regular – NoTask, Regular – TargetCounting, Irregular – NoTask, and Irregular – TargetCounting conditions, respectively.

The stimulation effect was defined as:

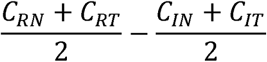

with positive values indicating higher connectivity during Regular stimulation. The task effect was defined as:

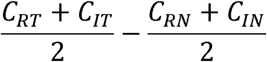

and the stimulation-by-task interaction as:

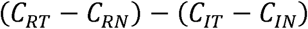

Regular–Irregular differences were also calculated separately for the NoTask and TargetCounting conditions.

To compare stimulation effects across region pairs, the 38-42 Hz stimulation effect was calculated separately for visual–parietal, visual-frontal, and parietal-frontal connectivity. The visual-parietal stimulation effect was compared separately with the visual–frontal and parietal–frontal effects.

To assess frequency specificity, connectivity for each region pair was additionally calculated for 4–7, 8–12, 15–30, 30–35, 38–42, and 44–48 Hz frequency bands. The upper band was restricted to 44–48 Hz to avoid the 49–51 Hz notch filter. The stimulation effect was calculated separately for each frequency band, and the 38–42 Hz effect was compared with the corresponding effect in each of the other five bands. The five comparisons were Holm-corrected separately for each region pair.

#### 2.5.3 Alpha-band connectivity

Alpha-band connectivity was analyzed to test whether task engagement was associated with connectivity changes in a frequency band related to visual attention and task demands. Connectivity for the three region pairs was calculated between 8 and 12 Hz using the same procedure described above. The PCC, sensor-space, and gamma–alpha analyses were performed for visual–parietal connectivity.

The primary alpha-band task effect was the TargetCounting–NoTask difference averaged across Regular and Irregular stimulation. The interaction was defined as the difference between the TargetCounting–NoTask contrasts under Regular and Irregular stimulation.

To compare condition effects between gamma and alpha connectivity, connectivity values were standardized separately for each band across all participant-condition observations. Stimulation and task effects were then calculated for each participant. Three within-participant contrasts were tested: the band-by-stimulation interaction (gamma minus alpha stimulation effect), the band-by-task interaction (gamma minus alpha task effect), and the band-by-stimulation-by-task interaction. Mean standardized gamma and alpha connectivity was also calculated for each of the four conditions.

Sensitivity analyses for alpha-band connectivity using PCC and sensor-space CSD are reported in Supplementary Table S2.

### 2.6 Robustness and control analyses

To test whether the results depended on the analysis pipeline, source-space connectivity was also estimated with a partial canonical coherence (PCC) beamformer in FieldTrip (release 20251218), with absolute imaginary coherence, and with Fourier-based dWPLI. Sensor-space connectivity was estimated after current source density (CSD) transformation, using occipital (O1/Oz/O2), parietal (P3/Pz/P4), and frontal (F3/Fz/F4) electrode groups that approximately overlie the source-space regions. To limit the number of comparisons, three electrodes per region were selected in advance (electrode pairs = 27). Parameters are given in Supplementary Table S3.

To reduce sensitivity to source leakage, symmetric orthogonalization of the ROI time series was performed and ipsilateral and interhemispheric ROI pairs were compared (Supplementary Material).

To estimate the contribution of activity phase-locked to the stimulus, the stimulus-locked component was removed before dWPLI was calculated. Because epochs were cut from continuous recordings and the flicker frequency, measured with the EEG sampling clock, differed slightly from 40 Hz, the stimulus phase drifted slowly across epochs and was not recorded directly. It was therefore estimated from the 40 Hz phase of the source-space visual ROIs using a constant frequency offset for each participant and condition. Epochs were aligned to this phase. Because this procedure requires the Fourier coefficients of each epoch, each 4-s epoch was multiplied by a Hanning window and Fourier transformed, the mean Fourier coefficient was subtracted, and dWPLI was calculated at 38–42 Hz from the remaining non-phase-locked coefficients. Details and alternative procedures are given in Supplementary Material.

### 2.7 Entrainment analyses

Source-space entrainment was measured in the visual, parietal, and frontal ROIs. Hanning-tapered Fourier coefficients were calculated at 40 Hz for each epoch. Inter-trial phase coherence (ITPC) was calculated across epochs and averaged across the ROIs of each region. Source power was calculated from the squared magnitude of the 40 Hz coefficients and log transformed.

As a check in sensor space, power was calculated from the mean signal across a predefined occipital electrode cluster (O1/Oz/O2/PO7/PO3/POz/PO4/PO8). Power between 39.5 and 40.5 Hz was averaged and log transformed. The 40 Hz signal-to-noise ratio was calculated as the Fourier amplitude at 40 Hz divided by the mean amplitude between 38 and 42 Hz, excluding 40 Hz (Khachatryan et al., 2022). Regular and Irregular stimulation were compared after averaging across the two task conditions.

### 2.8 Statistical analysis

Within-participant contrasts were tested using two-sided sign-flip permutation tests (100,000 permutations) and reported with Cohen’s d and bootstrap 95% confidence intervals. Cohen’s d was calculated from paired differences. Holm correction was applied to the five frequency-band comparisons and separately to the two comparisons across region pairs in source and sensor space. Holm correction was applied across the three region pairs when testing the stimulation effect. Participant-level bootstrap confidence intervals used 50,000 resamples. Edge-level maps were corrected using the Benjamini–Hochberg false discovery rate. To quantify evidence for the absence of a stimulation-by-task interaction (see section 2.5.2), Bayes factors (BF01) were computed from one-sample Bayesian t-tests on the within-participant interaction contrasts with a default Cauchy prior with scale 0.707 (Rouder et al., 2009). Statistical analyses used NumPy 2.5.0 and SciPy 1.18.0.

## 3. Results

### 3.1 Regular stimulation elicited 40 Hz entrainment

Compared with Irregular stimulation, Regular stimulation increased 40 Hz ITPC (*Cohen’s d* = 1.198) and log power (*Cohen’s d* = 1.882) in source-space visual regions. In parietal regions, 40 Hz ITPC and log power were also higher during Regular stimulation (Cohen’s d = 1.030 and 1.558), and smaller increases were observed in frontal regions (d = 0.580 and 0.412; Supplementary Table S4). Occipital sensor-space log power (*Cohen’s d* = 1.906) and SNR (Cohen’s d = 1.496) were also higher during Regular stimulation (all permutation *p* ≤ 2.0 × 10⁻□). The Regular-Irregular difference was positive in 28 of 29 participants for visual source-space ITPC and in all 29 participants for visual source-space power, sensor-space power, and sensor-space SNR. Values for each condition are reported in Supplementary Table S4.

### 3.2 Regular stimulation increased gamma-band connectivity

#### 3.2.1 Stimulation effects and control analyses

Regular stimulation increased source-space 38–42 Hz dWPLI for all three region pairs. The mean Regular-Irregular difference was 0.105 for visual–parietal connectivity (95% CI [0.076, 0.138], Cohen’s d = 1.216), with positive differences in all 29 participants (Fig. 3A–B). Visual–frontal connectivity increased by 0.020 (95% CI [0.013, 0.027], Cohen’s d = 1.080), with positive differences in 23 participants, and parietal–frontal connectivity increased by 0.023 (95% CI [0.014, 0.033], Cohen’s d = 0.856), with positive differences in 25 participants. All three effects remained significant after Holm correction (adjusted permutation p = 0.00003 for each comparison). Effects for individual source-space region pairs and sensor-space electrode pairs are shown in Fig. 4.

**Figure 3.**
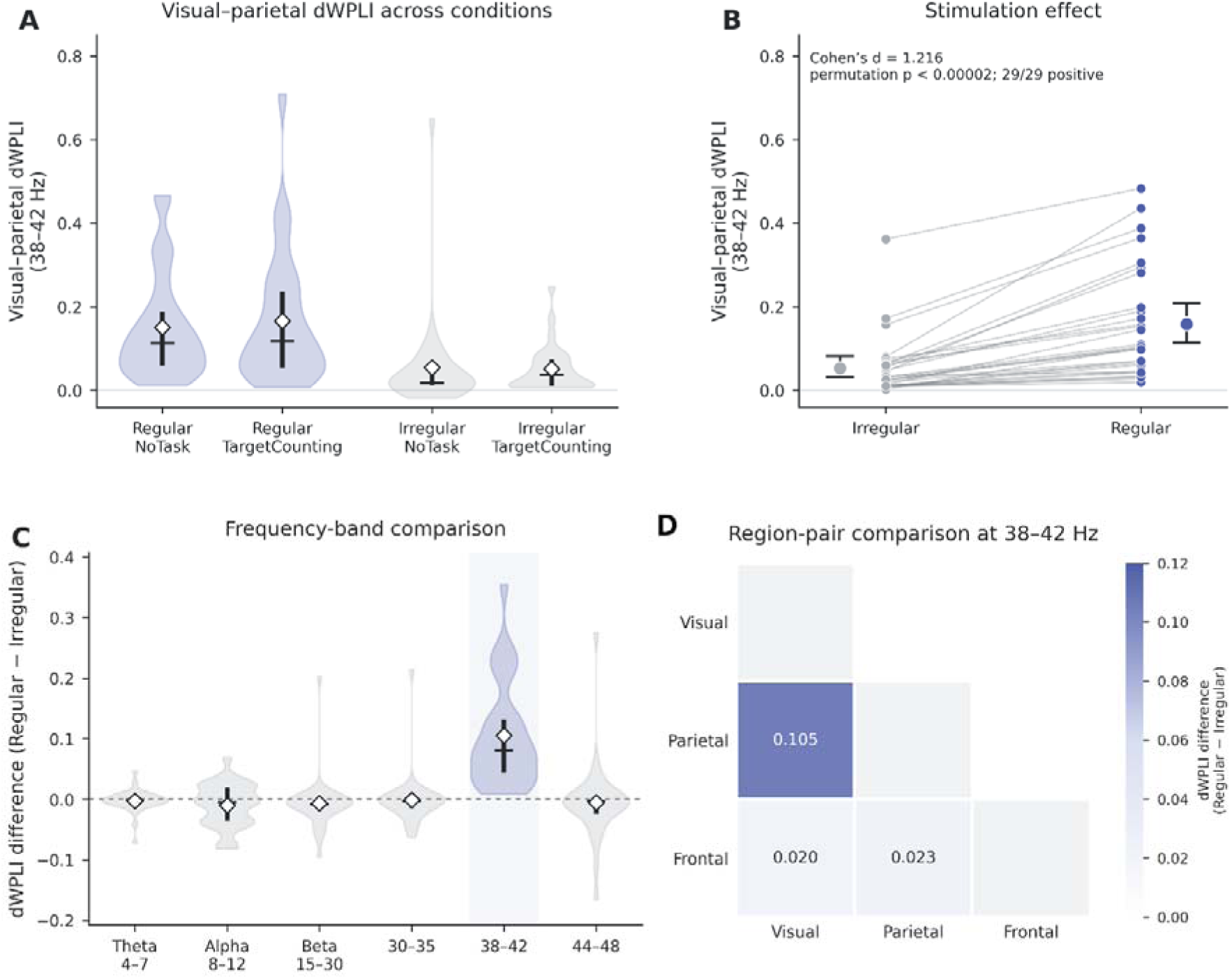
Regular stimulation increased visual–parietal gamma connectivity. **(A)** Source-space visual– parietal 38–42 Hz dWPLI across the four conditions. **(B)** Paired Irregular and Regular values after averaging across task conditions. Small points represent participants; lines connect paired observations; large points and error bars show the mean and 95% bootstrap confidence interval. **(C)** Visual–parietal stimulation effects across frequency bands. **(D)** Mean stimulation effects for the three region pairs. In A and C, violin width shows the estimated density, vertical lines the interquartile range, horizontal lines the median, and white diamonds the mean. All panels show source-space (wMNE) dWPLI.

**Figure 4.**
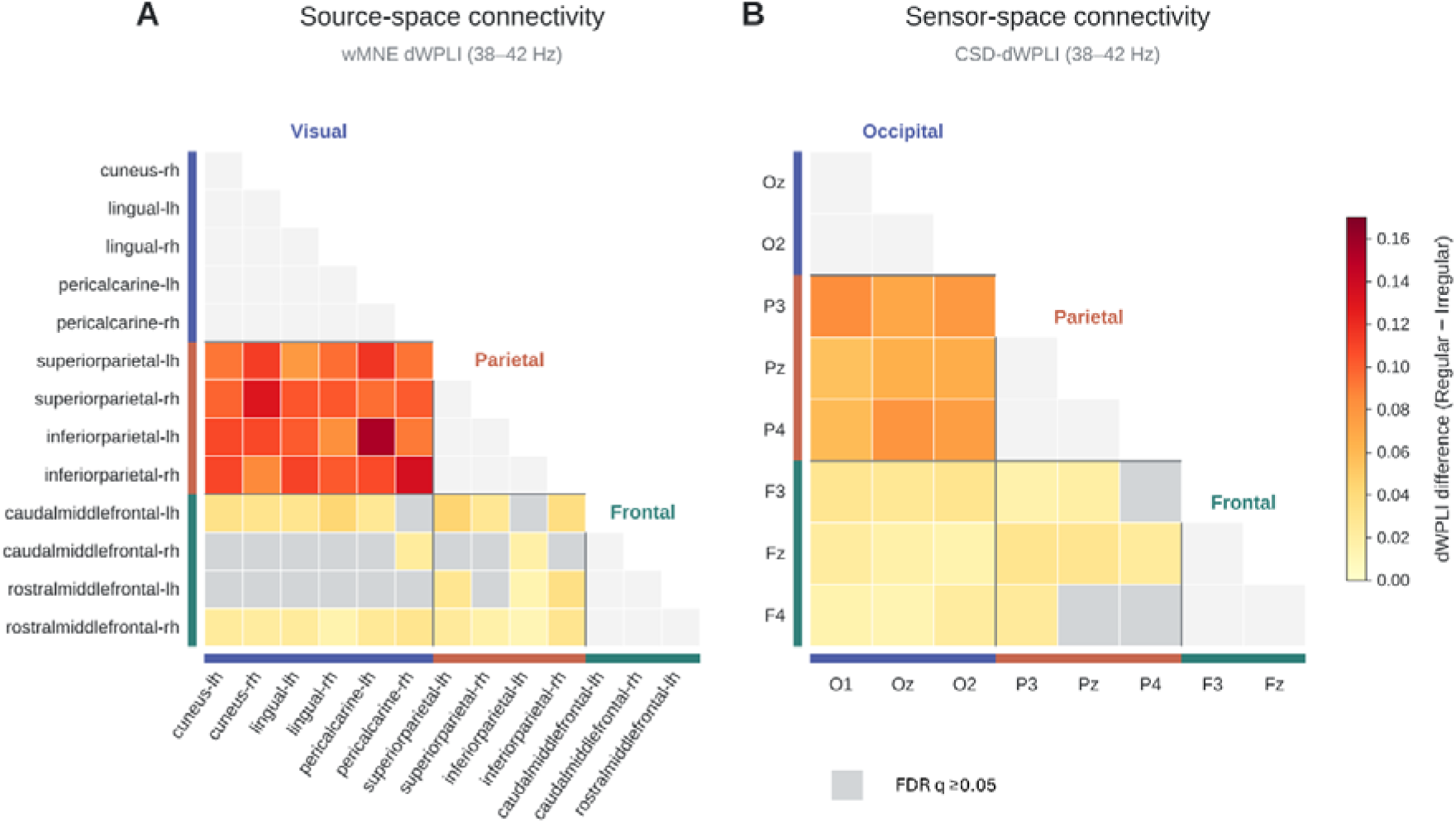
Changes in gamma connectivity during Regular versus Irregular stimulation. Mean Regular-Irregular differences in 38–42 Hz dWPLI are shown for **(A)** source-space region pairs estimated using wMNE and **(B)** sensor-space electrode pairs after CSD transformation. Differences were averaged across task conditions in 29 participants. Colored cells indicate significant increases (Benjamini–Hochberg FDR-corrected across 64 source-space connections or 27 electrode pairs).

The Regular–Irregular difference was significantly larger for visual–parietal connectivity compared to visual–frontal connectivity (Cohen’s d = 1.054, Holm-adjusted permutation p = 0.00002) and parietal–frontal connectivity (Cohen’s d = 1.024, Holm-adjusted permutation p = 0.00002).

The visual–parietal stimulation effect was also observed using alternative connectivity analyses (Table 1, Supplementary Table S5). PCC source reconstruction showed higher visual-parietal connectivity during Regular stimulation, and the sensor-space CSD analysis showed higher occipital–parietal connectivity, with positive differences in 28 participants.

**Table 1.** Visual–parietal 38–42 Hz connectivity effects across analysis methods.

| Analysis | Space | Measure | Cohen’s d | Permutation p | Positive participants |
|---|---|---|---|---|---|
| wMNE (PCA-flip) | Source | dWPLI | 1.216 | <0.00002 | 29/29 |
| PCC (all edges) | Source | dWPLI | 0.383 | 0.0148 | 19/29 |
| CSD | Sensor | dWPLI | 1.293 | <0.00002 | 28/29 |
| wMNE (PCA-flip) | Source | Absolute imaginary coherence | 1.033 | <0.00002 | 28/29 |

*Effects were calculated as Regular minus Irregular stimulation. Positive participants had a Regular-minus-Irregular value greater than zero. Permutation p values are two-sided (100,000 sign-flip permutations) and unadjusted.*

The visual–parietal effect remained after symmetric orthogonalization and when the analysis was restricted to interhemispheric ROI pairs. In contrast, the visual–frontal and parietal–frontal effects were reduced after orthogonalization (Supplementary Tables S6–S8).

Removal of the estimated stimulus-locked component substantially reduced the stimulation effects. Nevertheless, Regular stimulation increased visual–parietal connectivity (Cohen’s d = 0.48, Holm-adjusted permutation p = 0.011), with positive differences in 22 of 29 participants. The visual–frontal and parietal–frontal effects were no longer significant after removal of the stimulus-locked component (both Holm-adjusted p ≥ 0.52). Because stimulus phase was estimated from the source-space ROIs rather than recorded directly, the contribution of the stimulus-locked component is approximate; alternative procedures produced similar results (Supplementary Tables S9 and S10 and Fig. S1).

#### 3.2.2 Frequency specificity

The visual-parietal stimulation effect was largest around the stimulation frequency (Fig. 3C). The Regular-Irregular difference was larger at 38–42 Hz than in the theta, alpha, and beta bands and in the neighboring 30–35 and 44–48 Hz bands. Effect sizes for the five paired comparisons ranged from Cohen’s d = 0.910 to 1.353 (all Holm-adjusted permutation p < 0.0001). The same pattern was observed for visual– frontal and parietal–frontal connectivity (Cohen’s d = 0.449 to 0.962, all Holm-adjusted permutation p < 0.02; Supplementary Table S11).

As noted above, the magnitude of the 38–42 Hz stimulation effect differed across region pairs, with a larger effect for visual–parietal connectivity than for visual–frontal and parietal–frontal connectivity (Fig. 3D). The sensor-space CSD analysis showed the same pattern (Supplementary Table S5). In the PCC analysis, however, the visual–parietal effect did not differ from the visual–frontal or parietal–frontal effect (both Holm-adjusted p = 1.0).

### 3.3 Effects of the concurrent task on gamma connectivity

Regular stimulation increased source-space visual–parietal 38-42 Hz gamma connectivity under both conditions. The Regular-Irregular difference was 0.096 during NoTask (95% CI [0.044, 0.144], Cohen’s d = 0.690, Holm-adjusted permutation p = 0.002), with a positive difference in 26 of 29 participants, and 0.114 during TargetCounting (95% CI [0.074, 0.165], d = 0.884, Holm-adjusted p < 0.0001), with a positive difference in 28 participants.

However, the magnitude of the visual–parietal stimulation effect did not differ between NoTask and TargetCounting (interaction estimate = 0.018, 95% CI [−0.037, 0.103], Cohen’s d = 0.090, permutation p = 0.903), and the Bayes factor favored the absence of an interaction (BF01 = 4.6).

For visual–frontal and parietal–frontal connectivity, the Regular-Irregular differences were larger during TargetCounting (d = 0.623 and 0.410, both Holm-adjusted p < 0.0001) than during NoTask (d = 0.036 and 0.123, both Holm-adjusted p = 1.0). However, neither stimulation-by-task interaction was significant after Holm correction (d = 0.274 and 0.211; Holm-adjusted p = 0.098 and 0.639). The corresponding Bayes factors were inconclusive (BF01 = 1.9 and 2.8).

The stimulation-by-task interaction was also not significant in the PCC analysis (p = 0.48). In the sensor-space CSD analysis, the interaction reached p = 0.047 before correction (Cohen’s d = 0.31) but did not remain significant after Holm correction across the three analysis methods (Holm-adjusted p = 0.14). TargetCounting did not significantly change 40 Hz ITPC or power in source-space visual, parietal, or frontal regions (Supplementary Table S12).

### 3.4 Task effects on alpha connectivity

Averaged across stimulation conditions, visual–parietal alpha connectivity was higher during TargetCounting than NoTask in the wMNE analysis (Cohen’s d = 0.632, permutation p = 0.00135; Fig. 5A–B). The task effect was also significant in the PCC and the sensor-space CSD analysis (all p ≤ 0.0141). The task effect was not significant for visual–frontal (d = 0.214, p = 0.257) or parietal–frontal connectivity (d = 0.130, p = 0.495; Supplementary Table S2).

**Figure 5.**
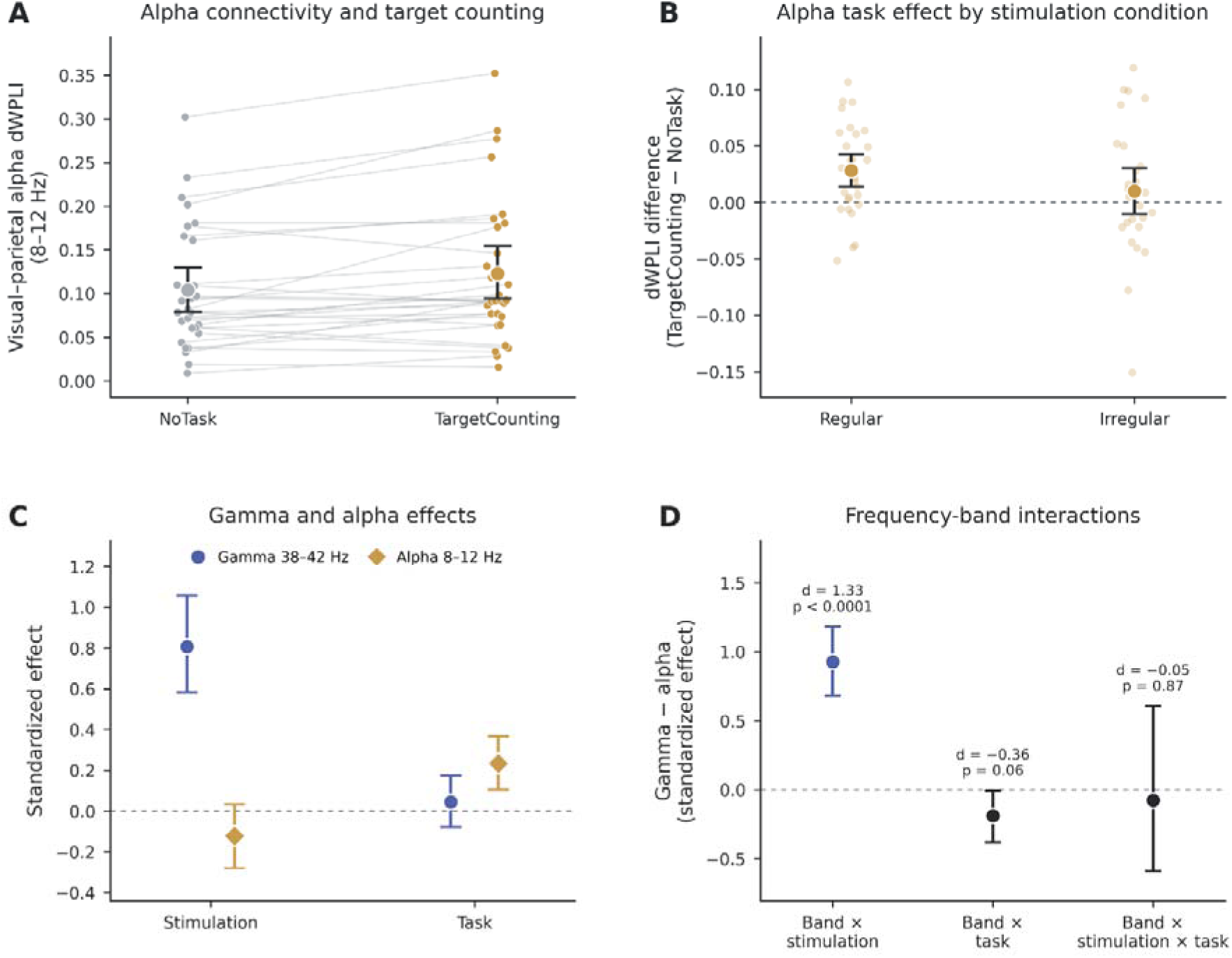
Alpha connectivity during target counting and comparison with gamma connectivity. **(A)** Visual– parietal alpha (8–12 Hz) dWPLI during NoTask and TargetCounting, averaged across stimulation conditions. Lines connect observations from the same participant. **(B)** TargetCounting–NoTask alpha effects under Regular and Irregular stimulation. Small points represent participants. **(C)** Standardized stimulation and task effects for visual–parietal gamma (38–42 Hz) and alpha connectivity. **(D)** Gamma-minus-alpha contrasts: band-by-stimulation, band-by-task, and band-by-stimulation-by-task interactions, with Cohen’s d and permutation p values. Large symbols show means; error bars show participant-bootstrap 95% confidence intervals. All panels show source-space (wMNE) dWPLI.

While source-space visual-parietal 38-42 Hz gamma connectivity showed a strong stimulation effect (Cohen’s d = 1.216, permutation p < 0.00002; Fig. 5C), its task effect was not significant (d = 0.127, p = 0.505). Alpha connectivity showed the opposite pattern; it increased during TargetCounting, but its stimulation effect was not significant (d = −0.275, p = 0.149).

After standardization, the stimulation effect was larger for gamma than for alpha connectivity (band-by-stimulation interaction: Cohen’s d = 1.326, permutation p < 0.00002), with a positive gamma-minus-alpha difference in 29 of 29 participants. The task effect did not differ significantly between the two bands (band-by-task interaction: d = −0.362, p = 0.062), and the three-way interaction was also not significant (p = 0.869; Fig. 5D).

## 4. Discussion

This study examined whether 40 Hz visual stimulation increases gamma-band connectivity between visual and higher-order cortical regions and whether this effect is modulated by a concurrent task. Regular stimulation increased 38–42 Hz connectivity between visual, parietal, and frontal regions, with the strongest and most robust increase between visual and parietal regions. The visual–parietal increase remained after correction for source leakage and removal of the stimulus-locked component, whereas the increases involving frontal regions did not. These findings suggest that, beyond a widespread stimulus-locked response, 40 Hz stimulation increases synchronization most clearly between visual and parietal regions. The concurrent task increased visual–parietal alpha connectivity but did not reliably modulate the gamma-connectivity effect, suggesting that stimulation and task demands affected this connection mainly in different frequency bands.

### 4.1 Gamma connectivity between visual, parietal and frontal regions

Regular stimulation increased 40 Hz power and ITPC in visual source space, as well as 40 Hz power and signal-to-noise ratio (SNR) at occipital sensors. These effects were consistent across nearly all participants and agree with previous human studies showing robust neural responses to 40 Hz sensory stimulation (Jones et al., 2019; Khachatryan et al., 2022; Longo et al., 2026; Mlinarič et al., 2026).

Regular stimulation also increased 38-42 Hz connectivity between visual, parietal, and frontal regions in the wMNE analysis, but the magnitude of this effect differed across region pairs. The increase was largest for visual–parietal connectivity. Previous studies found that gamma connectivity between the visual cortex and the inferior parietal lobule and precentral gyrus depended on the stimulation frequency (Petro et al., 2024) and that gamma connectivity was stronger for connections involving the visual cortex (Yoon et al., 2025). The present results extend these findings by showing that the visual–parietal effect remained after removal of the stimulus-locked component, whereas the increases in connectivity with frontal regions did not.

The larger visual–parietal increase was also observed in the sensor-space CSD analysis and with source-space absolute imaginary coherence, but not with PCC. The increase in visual–parietal connectivity remained after symmetric orthogonalization of the ROI time series and for interhemispheric ROI pairs. These analyses make it less likely that the effect can be explained entirely by source leakage, although residual source mixing cannot be excluded. In contrast, the visual–frontal and parietal–frontal effects were substantially reduced after orthogonalization and were no longer significant after removal of the stimulus-locked component.

The relative robustness of the visual–parietal effect is consistent with the close functional relationship between visual and parietal regions during visual processing and attentional selection (Corbetta and Shulman, 2002; Siegel et al., 2008; Vossel et al., 2014). However, increased 40 Hz power and SNR during Regular stimulation may also have improved phase estimation and thereby increased phase-based connectivity measures such as dWPLI (Schoffelen and Gross, 2009; Vinck et al., 2011). This alternative explanation applies to all region pairs and cannot be fully excluded with the present design.

The stimulation effect also showed clear frequency specificity. The Regular–Irregular difference in visual-parietal connectivity was larger at 38–42 Hz than in theta, alpha, beta, and the neighboring 30–35 Hz, and 44–48 Hz bands. This is consistent with previous SSVEP studies showing that neural responses to periodic visual stimulation are concentrated around the driving frequency (Herrmann, 2001; Srinivasan et al., 2006; Norcia et al., 2015).

Steady-state stimulation can drive multiple regions at the same frequency, so increased connectivity during stimulation may partly reflect stimulus-locked activity. Stimulus-locked activity can propagate from posterior regions with systematic phase delays (Tsoneva et al., 2021), which would produce consistent phase lags between regions. After removal of the estimated stimulus-locked component, the increases in visual–frontal and parietal–frontal connectivity were strongly reduced and no longer significant and therefore appear to be largely attributable to stimulus-locked activity. The remaining visual–parietal increase may reflect non-phase-locked synchronization, but it could also result from errors in the estimated stimulus phase or from nonstationarity of the response. Conversely, stimulus-locked activity is not limited to evoked responses: if the stimulation entrains endogenous oscillations, these would also be phase-locked to the stimulus and would be removed by this procedure. Distinguishing a superposition of evoked responses from entrained endogenous oscillations remains an unresolved issue in steady-state stimulation research (Duecker et al., 2021; Longo et al., 2026). Future studies could examine this distinction by directly recording the stimulus signal, testing for sustained oscillatory activity after stimulus offset, and comparing responses across different stimulation frequencies.

### 4.2 Task effects on gamma and alpha connectivity

We also examined whether TargetCounting modulated gamma connectivity associated with 40 Hz stimulation. The Regular-Irregular increase in visual–parietal gamma connectivity was observed during both NoTask and TargetCounting conditions, but the stimulation-by-task interaction was not significant. This result was consistent across the wMNE analysis and the PCC and sensor-space CSD analyses. The Bayes factor favored the absence of an interaction for visual–parietal connectivity.

Previous studies have reported mixed effects of task demands on 40 Hz responses, depending on the direction of attention and specific task requirements(Khachatryan et al., 2022; Ke et al., 2025; Mlinarič et al., 2025). These studies mainly examined local entrainment, scalp distribution, or intracranial phase relationships. In contrast, the present study examined task effects on connectivity between regions. In the present data, TargetCounting did not modify the visual–parietal stimulation effect.

For visual–frontal and parietal–frontal connectivity, the stimulation-by-task interaction was not significant after Holm correction, and the Bayes factors were inconclusive. The present data therefore neither support nor rule out a task effect for these region pairs. Descriptively, the stimulation effects on these region pairs were larger during TargetCounting than during NoTask, which is consistent in direction with the task-related spread of 40 Hz entrainment reported by Khachatryan et al. (2022). This pattern was not explained by a stronger 40 Hz response in visual regions, because 40 Hz ITPC and power in visual regions did not differ between the task conditions (Supplementary Table S12). One possibility is that the task engaged frontal regions through top-down attention, so that frontal activity followed the 40 Hz visual activity more closely (Siegel et al., 2008). However, the effects on these region pairs were not significant after removal of the stimulus-locked component. A larger sample is needed to test whether a task increases the spread of 40 Hz activity to frontal regions.

In contrast to gamma connectivity, visual-parietal alpha connectivity was higher during TargetCounting than during NoTask across the source-space (wMNE and PCC), and sensor-space (CSD) analyses. Alpha activity has been associated with visual selection, attentional allocation, and information exchange between visual and parietal regions (Jensen and Mazaheri, 2010; Foxe and Snyder, 2011; Marshall et al., 2015; Lobier et al., 2018). The present alpha-band effect therefore provides evidence that TargetCounting modulated visual-parietal connectivity. However, the alpha effect cannot be attributed uniquely to attention, because the task also involved working memory, and possibly differences in arousal and does not identify a specific cognitive process.

The direct comparison between frequency bands further showed that the stimulation regularity had a stronger effect on gamma than on alpha connectivity. In contrast, although the task increased alpha but not gamma connectivity when the bands were examined separately, the band-by-task interaction was not significant. Differences between alpha and gamma activity have also been reported in attention studies. Alpha activity has been found to vary mainly with prestimulus attentional state, while gamma activity is more closely related to subsequent stimulus processing (Bauer et al., 2014; van Ede et al., 2014). Scheeringa et al. found that alpha/beta and gamma activity separately explained BOLD changes during a visual attention task (Scheeringa et al., 2011). However, the two frequency bands are not completely independent, as pre-stimulus alpha activity can predict subsequent visually induced gamma activity (Popov et al., 2017). Together, these findings suggest that, for visual–parietal connectivity, stimulation regularity affected gamma more than alpha connectivity, whereas the evidence for a band-specific task effect was weaker.

### 4.3 Limitations

This study has several limitations. First, we included 29 healthy young adults and examined EEG changes during short-term stimulation. Whether these findings can be generalized to older adults and clinical populations, or whether they are associated with long-term effects or cognitive benefits, remains to be tested. Second, the design included only fixed 40 Hz and irregular flicker and a simple target-counting task. Our conclusions are therefore limited to the comparison between regular and irregular stimulation, and the absence of a reliable stimulation-by-task interaction does not exclude effects of other tasks or higher task loads. Third, EEG functional connectivity reflects statistical relationships between signals and cannot determine the direction of interactions. Source reconstruction used a template head model and standard electrode coordinates, and the analysis was limited to selected visual, parietal, and frontal regions. Stimulus timing was not recorded directly, so the stimulus phase was estimated from the EEG and the contribution of stimulus-locked activity is approximate. Finally, the robustness analyses were based on the same EEG dataset and do not replace replication in independent samples.

## Conclusion

This study showed that 40 Hz visual stimulation increased gamma-band connectivity between the visual cortex and higher-order regions, with the most robust increase between visual and parietal regions. Smaller visual–frontal and parietal–frontal effects did not remain significant after removal of the estimated stimulus-locked component. The concurrent target-counting task did not reliably modulate the visual– parietal effect, although it increased visual-parietal alpha connectivity. These findings highlight the importance of distinguishing stimulus-locked responses from interregional synchronization when using 40 Hz responses or connectivity measures to characterize the network effects of sensory stimulation. Future studies that record stimulus timing directly and use intracranial or high-density recordings could test more directly whether 40 Hz stimulation engages endogenous gamma synchronization in frontal and hippocampal networks, and whether these effects differ in older adults and patients with Alzheimer’s disease.

## Data Availability

The data supporting the findings of this study are available from the corresponding author upon reasonable request.

## Reference

Adaikkan, C., Middleton, S.J., Marco, A., Pao, P.-C., Mathys, H., Kim, D.N.-W., Gao, F., Young, J.Z., Suk, H.-J., Boyden, E.S., McHugh, T.J., Tsai, L.-H., 2019. Gamma Entrainment Binds Higher-Order Brain Regions and Offers Neuroprotection. Neuron 102, 929–943.e8. 10.1016/j.neuron.2019.04.011

Bastos, A.M., Schoffelen, J.-M., 2016. A Tutorial Review of Functional Connectivity Analysis Methods and Their Interpretational Pitfalls. Front. Syst. Neurosci. 9. 10.3389/fnsys.2015.00175

Bauer, M., Stenner, M.-P., Friston, K.J., Dolan, R.J., 2014. Attentional modulation of alpha/beta and gamma oscillations reflect functionally distinct processes. J. Neurosci. Off. J. Soc. Neurosci. 34, 16117–16125. 10.1523/JNEUROSCI.3474-13.2014

Chan, D., de Weck, G., Jackson, B.L., Suk, H.-J., Milman, N.P., Kitchener, E., Avalos, V.S.F., Quay, M.J., Aoki, K., Ruiz, E., Becker, A., Zheng, M., Philips, R., Firenze, R., Geigenmüller, U., Hammerschlag, B., Arnold, S., Kivisäkk, P., Brickhouse, M., Touroutoglou, A., Brown, E.N., Boyden, E.S., Dickerson, B.C., Klerman, E.B., Tsai, L.-H., 2025. Gamma sensory stimulation in mild Alzheimer’s dementia: An open-label extension study. Alzheimers Dement. J. Alzheimers Assoc. 21, e70792. 10.1002/alz.70792

Chan, D., Suk, H.-J., Jackson, B.L., Milman, N.P., Stark, D., Klerman, E.B., Kitchener, E., Avalos, V.S.F., Weck, G. de, Banerjee, A., Beach, S.D., Blanchard, J., Stearns, C., Boes, A.D., Uitermarkt, B., Gander, P., Iii, M.H., Sternberg, E.J., Nieto-Castanon, A., Anteraper, S., Whitfield-Gabrieli, S., Brown, E.N., Boyden, E.S., Dickerson, B.C., Tsai, L.-H., 2022. Gamma frequency sensory stimulation in mild probable Alzheimer’s dementia patients: Results of feasibility and pilot studies. PLOS ONE 17, e0278412. 10.1371/journal.pone.0278412

Corbetta, M., Shulman, G.L., 2002. Control of goal-directed and stimulus-driven attention in the brain. Nat. Rev. Neurosci. 3, 201–215. 10.1038/nrn755

Delorme, A., Makeig, S., 2004. EEGLAB: an open source toolbox for analysis of single-trial EEG dynamics including independent component analysis. J. Neurosci. Methods 134, 9–21. 10.1016/j.jneumeth.2003.10.009

Desikan, R.S., Ségonne, F., Fischl, B., Quinn, B.T., Dickerson, B.C., Blacker, D., Buckner, R.L., Dale, A.M., Maguire, R.P., Hyman, B.T., Albert, M.S., Killiany, R.J., 2006. An automated labeling system for subdividing the human cerebral cortex on MRI scans into gyral based regions of interest. NeuroImage 31, 968–980. 10.1016/j.neuroimage.2006.01.021

Duecker, K., Gutteling, T.P., Herrmann, C.S., Jensen, O., 2021. No Evidence for Entrainment: Endogenous Gamma Oscillations and Rhythmic Flicker Responses Coexist in Visual Cortex. J. Neurosci. 41, 6684–6698. 10.1523/JNEUROSCI.3134-20.2021

Farahibozorg, S.-R., Henson, R.N., Hauk, O., 2018. Adaptive cortical parcellations for source reconstructed EEG/MEG connectomes. NeuroImage 169, 23–45. 10.1016/j.neuroimage.2017.09.009

Fischl, B., 2012. FreeSurfer. NeuroImage, 20 YEARS OF fMRI 62, 774–781. 10.1016/j.neuroimage.2012.01.021

Foxe, J.J., Snyder, A.C., 2011. The Role of Alpha-Band Brain Oscillations as a Sensory Suppression Mechanism during Selective Attention. Front. Psychol. 2. 10.3389/fpsyg.2011.00154

Gramfort, A., Luessi, M., Larson, E., Engemann, D.A., Strohmeier, D., Brodbeck, C., Goj, R., Jas, M., Brooks, T., Parkkonen, L., Hämäläinen, M., 2013. MEG and EEG data analysis with MNE-Python. Front. Neurosci. 7. 10.3389/fnins.2013.00267

Herrmann, C.S., 2001. Human EEG responses to 1–100 Hz flicker: resonance phenomena in visual cortex and their potential correlation to cognitive phenomena. Exp. Brain Res. 137, 346–353. 10.1007/s002210100682

Iaccarino, H.F., Singer, A.C., Martorell, A.J., Rudenko, A., Gao, F., Gillingham, T.Z., Mathys, H., Seo, J., Kritskiy, O., Abdurrob, F., Adaikkan, C., Canter, R.G., Rueda, R., Brown, E.N., Boyden, E.S., Tsai, L.-H., 2016. Gamma frequency entrainment attenuates amyloid load and modifies microglia. Nature 540, 230–235. 10.1038/nature20587

Jensen, O., Mazaheri, A., 2010. Shaping Functional Architecture by Oscillatory Alpha Activity: Gating by Inhibition. Front. Hum. Neurosci. 4. 10.3389/fnhum.2010.00186

Jones, M., McDermott, B., Oliveira, B.L., O’Brien, A., Coogan, D., Lang, M., Moriarty, N., Dowd, E., Quinlan, L., Mc Ginley, B., Dunne, E., Newell, D., Porter, E., Elahi, M.A., O’ Halloran, M., Shahzad, A., 2019. Gamma Band Light Stimulation in Human Case Studies: Groundwork for Potential Alzheimer’s Disease Treatment. J. Alzheimer’s Dis. 70, 171–185. 10.3233/JAD-190299

Ke, S.-C., Lo, Y.-H., Tseng, P., 2025. The bidirectional effect of attention on gamma sensory stimulation: 40 Hz entrainment is weakened by externally-induced distraction but enhanced by internally-induced distraction. J. Alzheimers Dis. JAD 108, 353–366. 10.1177/13872877251376383

Khachatryan, E., Wittevrongel, B., Reinartz, M., Dauwe, I., Carrette, E., Meurs, A., Van Roost, D., Boon, P., Van Hulle, M.M., 2022. Cognitive tasks propagate the neural entrainment in response to a visual 40 Hz stimulation in humans. Front. Aging Neurosci. 14. 10.3389/fnagi.2022.1010765

Lobier, M., Palva, J.M., Palva, S., 2018. High-alpha band synchronization across frontal, parietal and visual cortex mediates behavioral and neuronal effects of visuospatial attention. NeuroImage 165, 222–237. 10.1016/j.neuroimage.2017.10.044

Longo, A.R., Bolognini, N., Zapparoli, L., 2026. Shaping gamma oscillations through sensory stimulation: A systematic review in healthy adults. Biol. Psychol. 204, 109211. 10.1016/j.biopsycho.2026.109211

Marshall, T.R., O’Shea, J., Jensen, O., Bergmann, T.O., 2015. Frontal Eye Fields Control Attentional Modulation of Alpha and Gamma Oscillations in Contralateral Occipitoparietal Cortex. J. Neurosci. 35, 1638–1647. 10.1523/JNEUROSCI.3116-14.2015

Martorell, A.J., Paulson, A.L., Suk, H.-J., Abdurrob, F., Drummond, G.T., Guan, W., Young, J.Z., Kim, D.N.-W., Kritskiy, O., Barker, S.J., Mangena, V., Prince, S.M., Brown, E.N., Chung, K., Boyden, E.S., Singer, A.C., Tsai, L.-H., 2019. Multi-sensory Gamma Stimulation Ameliorates Alzheimer’s-Associated Pathology and Improves Cognition. Cell 177, 256–271.e22. 10.1016/j.cell.2019.02.014

Mlinarič, T., Khachatryan, E., Wittevrongel, B., Dauwe, I., Carrette, E., Van Roost, D., Achten, E., Boon, P., Meurs, A., Van Hulle, M.M., 2025. Visual gamma stimulation induces 40 Hz neural oscillations in the human hippocampus and alters phase synchrony and lag. Commun. Biol. 8, 1301. 10.1038/s42003-025-08766-6

Mlinarič, T., Spruyt, L., Khachatryan, E., Wittevrongel, B., Reinartz, M., Van Laere, K., Dupont, P., Vandenberghe, R., Van Hulle, M.M., 2026. Early aperiodic EEG changes in preclinical and prodromal Alzheimer’s disease. Alzheimers Res. Ther. 18, 28. 10.1186/s13195-026-01953-5

Morgan, S.T., Hansen, J.C., Hillyard, S.A., 1996. Selective attention to stimulus location modulates the steady-state visual evoked potential. Proc. Natl. Acad. Sci. 93, 4770–4774. 10.1073/pnas.93.10.4770

Murdock, M.H., Yang, C.-Y., Sun, N., Pao, P.-C., Blanco-Duque, C., Kahn, M.C., Kim, T., Lavoie, N.S., Victor, M.B., Islam, M.R., Galiana, F., Leary, N., Wang, S., Bubnys, A., Ma, E., Akay, L.A., Sneve, M., Qian, Y., Lai, C., McCarthy, M.M., Kopell, N., Kellis, M., Piatkevich, K.D., Boyden, E.S., Tsai, L.-H., 2024. Multisensory gamma stimulation promotes glymphatic clearance of amyloid. Nature 627, 149–156. 10.1038/s41586-024-07132-6

Norcia, A.M., Appelbaum, L.G., Ales, J.M., Cottereau, B.R., Rossion, B., 2015. The steady-state visual evoked potential in vision research: A review. J. Vis. 15, 4. 10.1167/15.6.4

Petro, N.M., Webert, L.K., Springer, S.D., Okelberry, H.J., John, J.A., Horne, L.K., Glesinger, R., Rempe, M.P., Wilson, T.W., 2024. Optimal gamma-band entrainment of visual cortex. Hum. Brain Mapp. 45, e26775. 10.1002/hbm.26775

Popov, T., Kastner, S., Jensen, O., 2017. FEF-Controlled Alpha Delay Activity Precedes Stimulus-Induced Gamma-Band Activity in Visual Cortex. J. Neurosci. 37, 4117–4127. 10.1523/JNEUROSCI.3015-16.2017

Rouder, J.N., Speckman, P.L., Sun, D., Morey, R.D., Iverson, G., 2009. Bayesian t tests for accepting and rejecting the null hypothesis. Psychon. Bull. Rev. 16, 225–237. 10.3758/PBR.16.2.225

Scheeringa, R., Fries, P., Petersson, K.-M., Oostenveld, R., Grothe, I., Norris, D.G., Hagoort, P., Bastiaansen, M.C.M., 2011. Neuronal dynamics underlying high- and low-frequency EEG oscillations contribute independently to the human BOLD signal. Neuron 69, 572–583. 10.1016/j.neuron.2010.11.044

Schoffelen, J., Gross, J., 2009. Source connectivity analysis with MEG and EEG. Hum. Brain Mapp. 30, 1857–1865. 10.1002/hbm.20745

Siegel, M., Donner, T.H., Oostenveld, R., Fries, P., Engel, A.K., 2008. Neuronal synchronization along the dorsal visual pathway reflects the focus of spatial attention. Neuron 60, 709–719. 10.1016/j.neuron.2008.09.010

Srinivasan, R., Bibi, F.A., Nunez, P.L., 2006. Steady-State Visual Evoked Potentials: Distributed Local Sources and Wave-Like Dynamics Are Sensitive to Flicker Frequency. Brain Topogr. 18, 167–187. 10.1007/s10548-006-0267-4

Tsoneva, T., Garcia-Molina, G., Desain, P., 2021. SSVEP phase synchronies and propagation during repetitive visual stimulation at high frequencies. Sci. Rep. 11, 4975. 10.1038/s41598-021-83795-9

van Ede, F., Szebényi, S., Maris, E., 2014. Attentional modulations of somatosensory alpha, beta and gamma oscillations dissociate between anticipation and stimulus processing. NeuroImage 97, 134–141. 10.1016/j.neuroimage.2014.04.047

Vinck, M., Oostenveld, R., van Wingerden, M., Battaglia, F., Pennartz, C.M.A., 2011. An improved index of phase-synchronization for electrophysiological data in the presence of volume-conduction, noise and sample-size bias. NeuroImage 55, 1548–1565. 10.1016/j.neuroimage.2011.01.055

Vossel, S., Geng, J.J., Fink, G.R., 2014. Dorsal and ventral attention systems: distinct neural circuits but collaborative roles. Neurosci. Rev. J. Bringing Neurobiol. Neurol. Psychiatry 20, 150–159. 10.1177/1073858413494269

Yoon, E., Park, Y., Kim, H.J., Park, J., Han, J.W., Woo, S.J., Yoo, S., Kim, K.W., 2025. Center frequency as optimal frequency of visual stimulation for spreading entrained gamma rhythms to other target brain regions in cognitively normal older adults. GeroScience 47, 4451–4464. 10.1007/s11357-025-01552-6

