## Supplementary Materials for "40 Hz visual stimulation increases gamma-band functional connectivity beyond the visual cortex"

Supplementary Material

### Supplementary Methods

**Table S1. Epochs retained per condition**

| **Condition** | **Retained (%), mean** | **Epochs, mean** |
| --- | --- | --- |
| Regular–NoTask | 99.5 | 74.7 |
| Regular–TargetCounting | 99.0 | 74.2 |
| Irregular–NoTask | 97.8 | 73.4 |
| Irregular–TargetCounting | 96.4 | 72.3 |

*4-s epochs; 75 epochs per condition before rejection. Friedman test across conditions, p = 0.050.*

**Table S2. Alpha-band (8–12 Hz) task effects across methods and region pairs**

| **Method** | **Contrast** | **Mean difference** | **Cohen’s d** | **p** | **Positive** |
| --- | --- | --- | --- | --- | --- |
| wMNE dWPLI | Task effect (TargetCounting − NoTask) | 0.019 | 0.63 | 0.001 | 19/29 |
| wMNE dWPLI | Task effect, Regular | 0.028 | 0.70 | 0.0009 | 22/29 |
| wMNE dWPLI | Task effect, Irregular | 0.010 | 0.18 | 0.342 | 17/29 |
| wMNE dWPLI | Stimulation-by-task interaction | 0.018 | 0.23 | 0.222 | 20/29 |
| wMNE dWPLI (visual–frontal) | Task effect (TargetCounting − NoTask) | 0.005 | 0.21 | 0.257 | 16/29 |
| wMNE dWPLI (visual–frontal) | Task effect, Regular | 0.013 | 0.46 | 0.016 | 22/29 |
| wMNE dWPLI (visual–frontal) | Task effect, Irregular | −0.002 | −0.06 | 0.751 | 15/29 |
| wMNE dWPLI (visual–frontal) | Stimulation-by-task interaction | 0.015 | 0.31 | 0.100 | 17/29 |
| wMNE dWPLI (parietal–frontal) | Task effect (TargetCounting − NoTask) | 0.004 | 0.13 | 0.495 | 15/29 |
| wMNE dWPLI (parietal–frontal) | Task effect, Regular | 0.007 | 0.26 | 0.165 | 16/29 |
| wMNE dWPLI (parietal–frontal) | Task effect, Irregular | 0.000 | −0.01 | 0.978 | 14/29 |
| wMNE dWPLI (parietal–frontal) | Stimulation-by-task interaction | 0.008 | 0.13 | 0.491 | 16/29 |
| PCC dWPLI | Task effect (TargetCounting − NoTask) | 0.010 | 0.59 | 0.003 | 20/29 |
| PCC dWPLI | Task effect, Regular | 0.015 | 0.61 | 0.002 | 22/29 |
| PCC dWPLI | Task effect, Irregular | 0.005 | 0.20 | 0.285 | 18/29 |
| PCC dWPLI | Stimulation-by-task interaction | 0.011 | 0.31 | 0.111 | 19/29 |
| Sensor CSD dWPLI | Task effect (TargetCounting − NoTask) | 0.019 | 0.49 | 0.014 | 22/29 |
| Sensor CSD dWPLI | Task effect, Regular | 0.040 | 0.75 | 0.0004 | 25/29 |
| Sensor CSD dWPLI | Task effect, Irregular | −0.002 | −0.03 | 0.877 | 14/29 |
| Sensor CSD dWPLI | Stimulation-by-task interaction | 0.042 | 0.45 | 0.023 | 19/29 |

*p values unadjusted. Visual–parietal connectivity unless stated otherwise.*

**Table S3. Parameters of the connectivity analyses**

| **Analysis** | **Signals and source model** | **Spectral estimate and measure** |
| --- | --- | --- |
| wMNE dWPLI (main) | MNE-Python 1.12.1, MNE-Connectivity 0.8.1; fsaverage, 3-layer BEM, ico-3; depth 0.8, loose 0.2, λ² = 1/9, normal orientation; 14 regions, pca_flip | Multitaper, 7 DPSS tapers, ±1 Hz smoothing, 4-s epochs, 0.25 Hz resolution; averaged over band |
| wMNE \|iCoh\| | wMNE dWPLI | Absolute imaginary part of coherency; multitaper as above |
| PCC beamformer | FieldTrip 20251218; standard BEM; template source model, Desikan–Killiany labels; standard_1020 electrodes projected to scalp; common filter from all conditions (equal epoch numbers; 40 Hz); λ = 10%, fixed orientation | Hanning FFT, 1-Hz bins 38–42 Hz; dWPLI between all source points of two regions, averaged |
| Sensor CSD | FieldTrip spline CSD; standard_1020 positions; electrode groups O1/Oz/O2, P3/Pz/P4, F3/Fz/F4 | Hanning FFT, 1-Hz bins 38–42 Hz; dWPLI between all electrode pairs of two groups, averaged |
| Fourier-based dWPLI | wMNE regional time series | Hanning-tapered DFT of each 4-s epoch; dWPLI from complex coefficients in band |
| Symmetric orthogonalization | wMNE regional time series, 36–44 Hz FIR (251 taps), orthogonalized across 14 regions (Colclough et al., 2015) | Fourier-based dWPLI and absolute coherence, 38–42 Hz |
| Stimulus-locked removal | wMNE regional time series; stimulus phase modeled by Eq. S1 (grid search of Δf, −0.1 to 0.1 Hz, step 0.0002 Hz, maximizing visual 40 Hz ITPC) | Mean of phase-aligned coefficients subtracted (Eq. S2); alternatives in Eq. S3; Fourier-based dWPLI, 38–42 Hz |

*All analyses at 38–42 Hz unless stated otherwise.*

#### Removal of the stimulus-locked component

Stimulus phase in epoch $k$ (onset $t_{k}$; separate offset $\varphi_{h}$ for each 2.5-min stimulation $h$ of the 5-min recording):

$\theta_{k} = 2\pi\Delta f t_{k} + \varphi_{h}$	(S1)

Phase alignment of the Fourier coefficients $X_{r,k}(f)$ of region $r$ and subtraction of the mean over the epochs of half $h$:

$Z_{r,k}(f) = X_{r,k}(f) e^{-i\theta_{k}}, Y_{r,k}(f) = Z_{r,k}(f) - \langle Z_{r}(f)\rangle_{h}$	(S2)

Where $X_{r,k}(f)$ is the Fourier coefficient of region $r$ in epoch $k$ at frequency $f$ (38-42 Hz), $\theta_{k}$ is the stimulus phase from Eq. S1, $Z_{r,k}\left( f \right)$ is the phase-aligned coefficient, $\left\langle Z_{r}\left( f \right) \right\rangle_{h}$ is its mean over the epochs of half $h$ (the stimulus-locked component), and $Y_{r,k}\left( f \right)$ is the remaining non-phase-locked coefficient from which dWPLI was calculated. Because all regions in an epoch are rotated by the same angle, the alignment itself does not change dWPLI.

Alternative: subtraction of the mean over $n$ = 3, 5 or 10 consecutive epochs, scaled for the loss of variance:

$R_{r,k}(f) = [X_{r,k}(f) - \langle X_{r}(f)\rangle_{n}] \sqrt{\frac{n}{n - 1}}$	(S3)

A second alternative estimated the phase of epoch k from the visual 40 Hz phase of the three preceding and three following epochs (epoch k excluded).

### Supplementary Results

**Table S4. 40 Hz responses in the four conditions**

| **Measure** | **RN** | **RO** | **IN** | **IO** | **Regular − Irregular [95% CI]** | **Cohen’s d** | **p** | **Positive** |
| --- | --- | --- | --- | --- | --- | --- | --- | --- |
| Source ITPC (visual regions) | 0.285 | 0.265 | 0.109 | 0.100 | 0.171 [0.124, 0.225] | 1.20 | < 0.0001 | 28/29 |
| Source log power (visual regions) | −36.72 | −36.76 | −38.59 | −38.53 | 1.82 [1.48, 2.17] | 1.88 | < 0.0001 | 29/29 |
| Source ITPC (parietal regions) | 0.260 | 0.259 | 0.106 | 0.109 | 0.152 [0.105, 0.211] | 1.03 | < 0.0001 | 27/29 |
| Source log power (parietal regions) | −37.34 | −37.29 | −38.82 | −38.80 | 1.50 [1.16, 1.85] | 1.56 | < 0.0001 | 29/29 |
| Source ITPC (frontal regions) | 0.133 | 0.146 | 0.102 | 0.099 | 0.039 [0.017, 0.065] | 0.58 | 0.001 | 18/29 |
| Source log power (frontal regions) | −38.30 | −38.28 | −38.47 | −38.41 | 0.15 [0.02, 0.28] | 0.41 | 0.034 | 18/29 |
| Sensor log power (occipital) | 4.05 | 4.01 | 2.37 | 2.46 | 1.62 [1.32, 1.92] | 1.91 | < 0.0001 | 29/29 |
| Sensor SNR (occipital) | 5.22 | 4.97 | 1.00 | 1.01 | 4.09 [3.13, 5.07] | 1.50 | < 0.0001 | 29/29 |

*RN/RO: Regular–NoTask/TargetCounting; IN/IO: Irregular–NoTask/TargetCounting. SNR: amplitude at 40 Hz divided by the mean amplitude at 38–42 Hz excluding 40 Hz.*

**Table S5. Stimulation effects and region-pair comparisons across methods**

| **Method** | **Contrast** | **Mean difference [95% CI]** | **Cohen’s d** | **p (Holm)** | **Positive** |
| --- | --- | --- | --- | --- | --- |
| wMNE dWPLI | Visual–parietal stimulation effect | 0.105 [0.076, 0.137] | 1.22 | < 0.0001 | 29/29 |
| wMNE dWPLI | Visual–frontal stimulation effect | 0.020 [0.013, 0.027] | 1.08 | < 0.0001 | 23/29 |
| wMNE dWPLI | Parietal–frontal stimulation effect | 0.023 [0.014, 0.033] | 0.86 | < 0.0001 | 25/29 |
| wMNE dWPLI | Visual–parietal vs. visual–frontal | 0.085 [0.058, 0.116] | 1.05 | < 0.0001 | 27/29 |
| wMNE dWPLI | Visual–parietal vs. parietal–frontal | 0.082 [0.055, 0.113] | 1.02 | < 0.0001 | 27/29 |
| PCC dWPLI | Visual–parietal stimulation effect | 0.006 [0.001, 0.012] | 0.38 | 0.016 | 19/29 |
| PCC dWPLI | Visual–frontal stimulation effect | 0.005 [0.002, 0.008] | 0.53 | 0.016 | 20/29 |
| PCC dWPLI | Parietal–frontal stimulation effect | 0.006 [0.003, 0.009] | 0.72 | 0.0004 | 22/29 |
| PCC dWPLI | Visual–parietal vs. visual–frontal | 0.001 [−0.004, 0.007] | 0.08 | 1.000 | 16/29 |
| PCC dWPLI | Visual–parietal vs. parietal–frontal | 0.000 [−0.004, 0.006] | 0.03 | 1.000 | 14/29 |
| Sensor CSD dWPLI | Visual–parietal stimulation effect | 0.071 [0.052, 0.091] | 1.29 | < 0.0001 | 28/29 |
| Sensor CSD dWPLI | Visual–frontal stimulation effect | 0.021 [0.015, 0.029] | 1.06 | < 0.0001 | 27/29 |
| Sensor CSD dWPLI | Parietal–frontal stimulation effect | 0.020 [0.012, 0.028] | 0.84 | < 0.0001 | 25/29 |
| Sensor CSD dWPLI | Visual–parietal vs. visual–frontal | 0.049 [0.033, 0.067] | 1.04 | < 0.0001 | 26/29 |
| Sensor CSD dWPLI | Visual–parietal vs. parietal–frontal | 0.051 [0.036, 0.067] | 1.14 | < 0.0001 | 26/29 |
| wMNE \|iCoh\| | Visual–parietal stimulation effect | 0.036 [0.024, 0.049] | 1.03 | < 0.0001 | 28/29 |
| wMNE \|iCoh\| | Visual–frontal stimulation effect | 0.009 [0.007, 0.011] | 1.56 | < 0.0001 | 27/29 |
| wMNE \|iCoh\| | Parietal–frontal stimulation effect | 0.009 [0.005, 0.015] | 0.64 | 0.0001 | 26/29 |
| wMNE \|iCoh\| | Visual–parietal vs. visual–frontal | 0.027 [0.016, 0.040] | 0.82 | < 0.0001 | 26/29 |
| wMNE \|iCoh\| | Visual–parietal vs. parietal–frontal | 0.026 [0.014, 0.039] | 0.73 | 0.0001 | 26/29 |

*Regular − Irregular, averaged across task conditions. Holm correction across the three region pairs (stimulation effects) or the two comparisons (region-pair comparisons) within each method.*

**Table S6. Stimulation effects after symmetric orthogonalization**

| **Measure** | **Region pair** | **Mean difference [95% CI]** | **Cohen’s d** | **p (Holm)** | **Positive** |
| --- | --- | --- | --- | --- | --- |
| Orthogonalized dWPLI | Visual–parietal | 0.0207 [0.0145, 0.0274] | 1.14 | < 0.0001 | 28/29 |
| Orthogonalized dWPLI | Visual–frontal | 0.0022 [0.0007, 0.0040] | 0.48 | 0.003 | 19/29 |
| Orthogonalized dWPLI | Parietal–frontal | 0.0016 [0.0006, 0.0027] | 0.52 | 0.005 | 17/29 |
| Orthogonalized absolute coherence | Visual–parietal | 0.0249 [0.0199, 0.0296] | 1.81 | < 0.0001 | 28/29 |
| Orthogonalized absolute coherence | Visual–frontal | 0.0017 [-0.0029, 0.0058] | 0.14 | 0.486 | 18/29 |
| Orthogonalized absolute coherence | Parietal–frontal | 0.0029 [-0.0008, 0.0066] | 0.28 | 0.276 | 18/29 |

*Holm correction across the three region pairs within each measure.*

**Table S7. Visual–parietal effect compared with the other region pairs after symmetric orthogonalization**

| **Measure** | **Comparison** | **Cohen’s d** | **p (Holm)** | **Positive** |
| --- | --- | --- | --- | --- |
| Orthogonalized dWPLI | Visual–parietal vs. visual–frontal | 0.99 | < 0.0001 | 27/29 |
| Orthogonalized dWPLI | Visual–parietal vs. parietal–frontal | 1.04 | < 0.0001 | 28/29 |
| Orthogonalized absolute coherence | Visual–parietal vs. visual–frontal | 1.23 | < 0.0001 | 27/29 |
| Orthogonalized absolute coherence | Visual–parietal vs. parietal–frontal | 1.21 | < 0.0001 | 25/29 |

*Holm correction across the two comparisons within each measure.*

**Table S8. Stimulation effects for ipsilateral and interhemispheric region pairs (wMNE dWPLI)**

| **Region pair** | **Pair type** | **N pairs** | **Mean difference** | **Cohen’s d** | **p** | **Positive** |
| --- | --- | --- | --- | --- | --- | --- |
| Visual–parietal | Ipsilateral | 12 | 0.1094 | 1.21 | < 0.0001 | 29/29 |
| Visual–parietal | Interhemispheric | 12 | 0.1010 | 1.15 | < 0.0001 | 29/29 |
| Visual–parietal | Ipsilateral minus interhemispheric |  | 0.0083 | 0.19 | 0.305 | 19/29 |
| Visual–frontal | Ipsilateral | 12 | 0.0194 | 0.95 | < 0.0001 | 24/29 |
| Visual–frontal | Interhemispheric | 12 | 0.0205 | 1.08 | < 0.0001 | 24/29 |
| Visual–frontal | Ipsilateral minus interhemispheric |  | -0.0010 | -0.07 | 0.703 | 12/29 |
| Parietal–frontal | Ipsilateral | 8 | 0.0235 | 0.85 | < 0.0001 | 25/29 |
| Parietal–frontal | Interhemispheric | 8 | 0.0220 | 0.75 | < 0.0001 | 26/29 |
| Parietal–frontal | Ipsilateral minus interhemispheric |  | 0.0016 | 0.08 | 0.689 | 13/29 |

*p values unadjusted.*

**Table S9. Stimulation effects before and after removal of the stimulus-locked component**

| **Procedure** | **Region pair** | **Mean difference [95% CI]** | **Cohen’s d** | **p (Holm)** | **Positive** | **% of original** |
| --- | --- | --- | --- | --- | --- | --- |
| No removal | Visual–parietal | 0.0724 [0.0409, 0.1095] | 0.75 | < 0.0001 | 27/29 | 100% |
| No removal | Visual–frontal | 0.0055 [0.0031, 0.0084] | 0.74 | < 0.0001 | 23/29 | 100% |
| No removal | Parietal–frontal | 0.0069 [0.0029, 0.0129] | 0.48 | 0.00010 | 23/29 | 100% |
| Average over all epochs removed | Visual–parietal | 0.0526 [0.0314, 0.0764] | 0.82 | < 0.0001 | 27/29 | 73% |
| Average over all epochs removed | Visual–frontal | 0.0035 [0.0019, 0.0053] | 0.74 | 0.00020 | 22/29 | 63% |
| Average over all epochs removed | Parietal–frontal | 0.0061 [0.0021, 0.0123] | 0.40 | 0.00021 | 22/29 | 88% |
| Linear phase drift model | Visual–parietal | 0.0108 [0.0037, 0.0195] | 0.48 | 0.011 | 22/29 | 15% |
| Linear phase drift model | Visual–frontal | 0.0002 [-0.0017, 0.0024] | 0.03 | 0.898 | 11/29 | 3% |
| Linear phase drift model | Parietal–frontal | 0.0036 [-0.0005, 0.0106] | 0.21 | 0.517 | 19/29 | 52% |
| Phase from neighboring epochs | Visual–parietal | 0.0123 [0.0050, 0.0210] | 0.54 | 0.002 | 22/29 | 17% |
| Phase from neighboring epochs | Visual–frontal | 0.0005 [-0.0012, 0.0025] | 0.10 | 0.650 | 14/29 | 9% |
| Phase from neighboring epochs | Parietal–frontal | 0.0039 [-0.0002, 0.0108] | 0.23 | 0.268 | 17/29 | 56% |
| Average over 3 epochs removed | Visual–parietal | 0.0102 [0.0024, 0.0192] | 0.43 | 0.074 | 19/29 | 14% |
| Average over 3 epochs removed | Visual–frontal | 0.0003 [-0.0023, 0.0032] | 0.04 | 1.000 | 14/29 | 5% |
| Average over 3 epochs removed | Parietal–frontal | 0.0035 [-0.0010, 0.0113] | 0.18 | 1.000 | 16/29 | 51% |
| Average over 5 epochs removed | Visual–parietal | 0.0100 [0.0027, 0.0186] | 0.45 | 0.041 | 19/29 | 14% |
| Average over 5 epochs removed | Visual–frontal | -0.0003 [-0.0025, 0.0020] | -0.05 | 0.828 | 16/29 | -6% |
| Average over 5 epochs removed | Parietal–frontal | 0.0033 [-0.0009, 0.0103] | 0.19 | 0.828 | 14/29 | 47% |
| Average over 10 epochs removed | Visual–parietal | 0.0127 [0.0052, 0.0216] | 0.55 | 0.002 | 18/29 | 18% |
| Average over 10 epochs removed | Visual–frontal | 0.0001 [-0.0019, 0.0024] | 0.02 | 0.924 | 14/29 | 2% |
| Average over 10 epochs removed | Parietal–frontal | 0.0034 [-0.0006, 0.0102] | 0.20 | 0.648 | 15/29 | 49% |

*Fourier-based dWPLI. Linear phase drift model: procedure of the main text. Estimated frequency offset (Regular conditions): median −0.0031 Hz (IQR −0.0044 to −0.0024). Percentages are unreliable when the original effect is small.*

**Table S10. Visual–parietal effect compared with the other region pairs before and after removal of the stimulus-locked component**

| **Procedure** | **Comparison** | **Cohen’s d** | **p (Holm)** | **Positive** |
| --- | --- | --- | --- | --- |
| No removal | Visual–parietal vs. visual–frontal | 0.70 | < 0.0001 | 28/29 |
| No removal | Visual–parietal vs. parietal–frontal | 0.68 | < 0.0001 | 26/29 |
| Average over all epochs removed | Visual–parietal vs. visual–frontal | 0.78 | < 0.0001 | 28/29 |
| Average over all epochs removed | Visual–parietal vs. parietal–frontal | 0.71 | < 0.0001 | 25/29 |
| Linear phase drift model | Visual–parietal vs. visual–frontal | 0.49 | 0.003 | 22/29 |
| Linear phase drift model | Visual–parietal vs. parietal–frontal | 0.30 | 0.121 | 18/29 |
| Phase from neighboring epochs | Visual–parietal vs. visual–frontal | 0.53 | 0.002 | 22/29 |
| Phase from neighboring epochs | Visual–parietal vs. parietal–frontal | 0.36 | 0.064 | 18/29 |
| Average over 3 epochs removed | Visual–parietal vs. visual–frontal | 0.44 | 0.037 | 19/29 |
| Average over 3 epochs removed | Visual–parietal vs. parietal–frontal | 0.30 | 0.123 | 20/29 |
| Average over 5 epochs removed | Visual–parietal vs. visual–frontal | 0.47 | 0.013 | 19/29 |
| Average over 5 epochs removed | Visual–parietal vs. parietal–frontal | 0.26 | 0.170 | 19/29 |
| Average over 10 epochs removed | Visual–parietal vs. visual–frontal | 0.54 | 0.002 | 21/29 |
| Average over 10 epochs removed | Visual–parietal vs. parietal–frontal | 0.34 | 0.077 | 21/29 |

*Holm correction across the two comparisons within each procedure.*

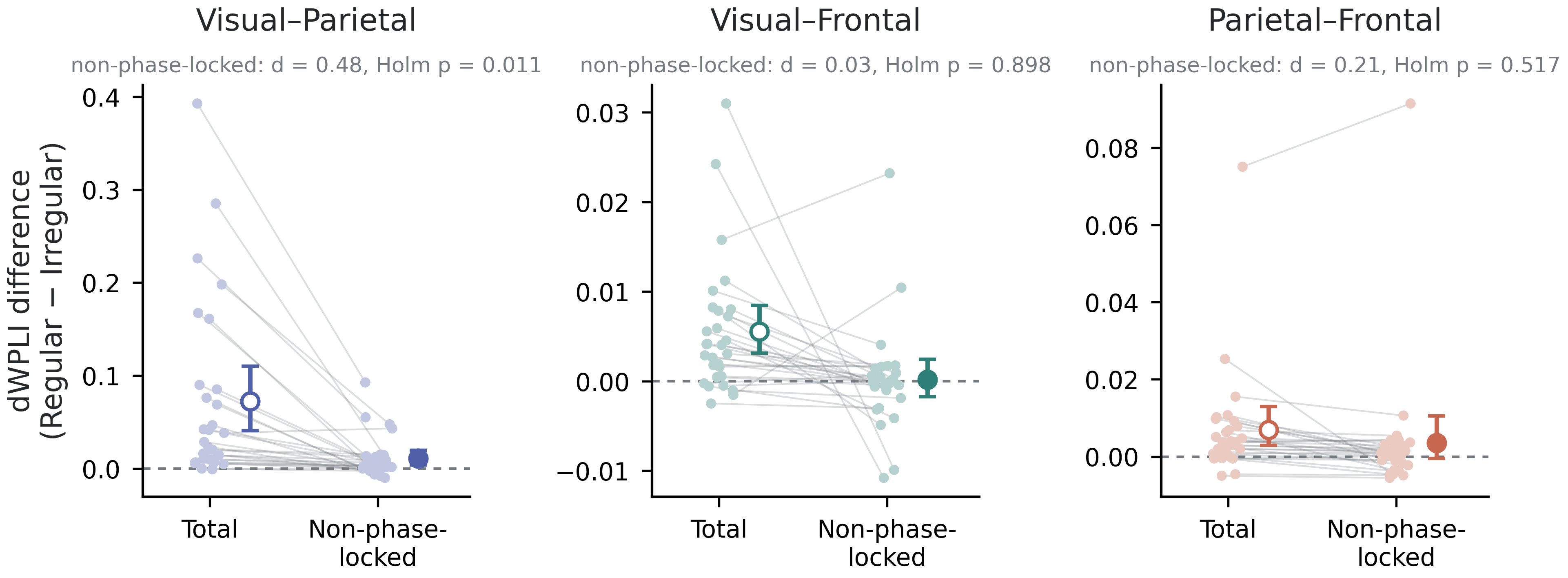

***Figure S1. Stimulation effects before and after removal of the stimulus-locked component****. Regular − Irregular 38–42 Hz Fourier-based dWPLI before (Total) and after removal (Non-phase-locked). Lines: participants; large symbols: mean and 95% bootstrap CI. Different y-axis scales.*

**Table S11. Frequency specificity of the stimulation effect for the three region pairs (wMNE dWPLI)**

| **Region pair** | **Contrast** | **Mean difference [95% CI]** | **Cohen’s d** | **p (Holm)** | **Positive** |
| --- | --- | --- | --- | --- | --- |
| Visual–parietal | Stimulation effect, 38–42 Hz | 0.105 [0.076, 0.137] | 1.22 | < 0.0001 | 29/29 |
| Visual–parietal | 38–42 Hz vs. theta (4–7 Hz) | 0.108 [0.079, 0.140] | 1.26 | < 0.0001 | 28/29 |
| Visual–parietal | 38–42 Hz vs. alpha (8–12 Hz) | 0.115 [0.086, 0.147] | 1.35 | < 0.0001 | 29/29 |
| Visual–parietal | 38–42 Hz vs. beta (15–30 Hz) | 0.112 [0.077, 0.149] | 1.13 | < 0.0001 | 28/29 |
| Visual–parietal | 38–42 Hz vs. 30–35 Hz | 0.106 [0.069, 0.145] | 1.01 | < 0.0001 | 27/29 |
| Visual–parietal | 38–42 Hz vs. 44–48 Hz | 0.111 [0.068, 0.155] | 0.91 | < 0.0001 | 27/29 |
| Visual–frontal | Stimulation effect, 38–42 Hz | 0.020 [0.013, 0.027] | 1.08 | < 0.0001 | 23/29 |
| Visual–frontal | 38–42 Hz vs. theta (4–7 Hz) | 0.018 [0.002, 0.032] | 0.45 | 0.016 | 24/29 |
| Visual–frontal | 38–42 Hz vs. alpha (8–12 Hz) | 0.024 [0.013, 0.036] | 0.75 | 0.0008 | 25/29 |
| Visual–frontal | 38–42 Hz vs. beta (15–30 Hz) | 0.021 [0.013, 0.029] | 0.91 | 0.0002 | 25/29 |
| Visual–frontal | 38–42 Hz vs. 30–35 Hz | 0.016 [0.005, 0.026] | 0.56 | 0.010 | 22/29 |
| Visual–frontal | 38–42 Hz vs. 44–48 Hz | 0.021 [0.008, 0.034] | 0.57 | 0.008 | 22/29 |
| Parietal–frontal | Stimulation effect, 38–42 Hz | 0.023 [0.014, 0.033] | 0.86 | < 0.0001 | 25/29 |
| Parietal–frontal | 38–42 Hz vs. theta (4–7 Hz) | 0.030 [0.019, 0.042] | 0.96 | < 0.0001 | 27/29 |
| Parietal–frontal | 38–42 Hz vs. alpha (8–12 Hz) | 0.034 [0.018, 0.054] | 0.67 | < 0.0001 | 25/29 |
| Parietal–frontal | 38–42 Hz vs. beta (15–30 Hz) | 0.021 [0.013, 0.029] | 0.96 | < 0.0001 | 26/29 |
| Parietal–frontal | 38–42 Hz vs. 30–35 Hz | 0.016 [0.008, 0.024] | 0.74 | 0.0009 | 24/29 |
| Parietal–frontal | 38–42 Hz vs. 44–48 Hz | 0.022 [0.010, 0.036] | 0.64 | 0.0009 | 24/29 |

*Regular − Irregular, averaged across task conditions. Stimulation effects at 38–42 Hz as in Table S5. Holm correction across the five frequency-band comparisons within each region pair.*

**Table S12. Task effects on the 40 Hz response in visual, parietal, and frontal regions**

| **Measure** | **Contrast** | **Mean difference [95% CI]** | **Cohen’s d** | **p (Holm)** |
| --- | --- | --- | --- | --- |
| ITPC (visual) | TargetCounting − NoTask (Regular) | −0.019 [−0.081, 0.040] | −0.11 | 1.00 |
| ITPC (visual) | TargetCounting − NoTask (Irregular) | −0.009 [−0.027, 0.009] | −0.17 | 1.00 |
| ITPC (visual) | Stimulation-by-task interaction | −0.011 [−0.079, 0.054] | −0.06 | 1.00 |
| Log power (visual) | TargetCounting − NoTask (Regular) | −0.039 [−0.129, 0.060] | −0.15 | 1.00 |
| Log power (visual) | TargetCounting − NoTask (Irregular) | 0.059 [−0.144, 0.288] | 0.10 | 1.00 |
| Log power (visual) | Stimulation-by-task interaction | −0.098 [−0.364, 0.142] | −0.14 | 1.00 |
| ITPC (parietal) | TargetCounting − NoTask (Regular) | −0.001 [−0.055, 0.050] | −0.01 | 1.00 |
| ITPC (parietal) | TargetCounting − NoTask (Irregular) | 0.003 [−0.014, 0.018] | 0.06 | 1.00 |
| ITPC (parietal) | Stimulation-by-task interaction | −0.003 [−0.060, 0.051] | −0.02 | 1.00 |
| Log power (parietal) | TargetCounting − NoTask (Regular) | 0.050 [−0.038, 0.137] | 0.21 | 0.83 |
| Log power (parietal) | TargetCounting − NoTask (Irregular) | 0.020 [−0.140, 0.185] | 0.04 | 1.00 |
| Log power (parietal) | Stimulation-by-task interaction | 0.030 [−0.161, 0.221] | 0.06 | 1.00 |
| ITPC (frontal) | TargetCounting − NoTask (Regular) | 0.013 [−0.008, 0.036] | 0.21 | 0.84 |
| ITPC (frontal) | TargetCounting − NoTask (Irregular) | −0.002 [−0.018, 0.013] | −0.05 | 0.84 |
| ITPC (frontal) | Stimulation-by-task interaction | 0.015 [−0.011, 0.041] | 0.20 | 0.84 |
| Log power (frontal) | TargetCounting − NoTask (Regular) | 0.017 [−0.151, 0.195] | 0.03 | 1.00 |
| Log power (frontal) | TargetCounting − NoTask (Irregular) | 0.061 [−0.157, 0.284] | 0.10 | 1.00 |
| Log power (frontal) | Stimulation-by-task interaction | −0.044 [−0.336, 0.258] | −0.05 | 1.00 |

*Holm correction across the three contrasts within each measure.*

**Table S13. Fourier-based dWPLI**

| **Contrast** | **Mean difference [95% CI]** | **Cohen’s d** | **p (Holm)** | **Positive** |
| --- | --- | --- | --- | --- |
| Stimulation effect, visual–parietal, 38–42 Hz | 0.072 [0.041, 0.110] | 0.75 | < 0.0001 | 27/29 |
| 38–42 Hz vs. theta (4–7 Hz) | 0.074 [0.043, 0.111] | 0.77 | < 0.0001 | 29/29 |
| 38–42 Hz vs. alpha (8–12 Hz) | 0.071 [0.040, 0.109] | 0.74 | < 0.0001 | 26/29 |
| 38–42 Hz vs. beta (15–30 Hz) | 0.071 [0.037, 0.110] | 0.69 | 0.0001 | 28/29 |
| 38–42 Hz vs. 30–35 Hz | 0.068 [0.032, 0.108] | 0.63 | 0.0006 | 27/29 |
| Visual–parietal vs. visual–frontal | 0.067 [0.036, 0.104] | 0.70 | < 0.0001 | 28/29 |
| Visual–parietal vs. parietal–frontal | 0.065 [0.034, 0.102] | 0.68 | < 0.0001 | 26/29 |

*Holm correction across the four frequency-band comparisons and across the two region-pair comparisons.*

**Table S14. Sensitivity analysis for one participant with a replaced block**

| **Data** | **Region pair** | **Stimulation d** | **p (Holm)** | **Positive** | **Interaction d** | **Interaction p (Holm)** |
| --- | --- | --- | --- | --- | --- | --- |
| All data (main analysis) | Visual–parietal | 1.22 | < 0.0001 | 29/29 | 0.09 | 0.905 |
| All data (main analysis) | Visual–frontal | 1.08 | < 0.0001 | 23/29 | 0.27 | 0.095 |
| All data (main analysis) | Parietal–frontal | 0.86 | < 0.0001 | 25/29 | 0.21 | 0.639 |
| Non-duplicated data only | Visual–parietal | 1.21 | < 0.0001 | 29/29 | 0.09 | 0.892 |
| Non-duplicated data only | Visual–frontal | 1.08 | < 0.0001 | 23/29 | 0.28 | 0.093 |
| Non-duplicated data only | Parietal–frontal | 0.83 | < 0.0001 | 25/29 | 0.22 | 0.579 |
| Participant excluded (N = 28) | Visual–parietal | 1.21 | < 0.0001 | 28/28 | 0.09 | 0.903 |
| Participant excluded (N = 28) | Visual–frontal | 1.05 | < 0.0001 | 22/28 | 0.26 | 0.177 |
| Participant excluded (N = 28) | Parietal–frontal | 0.85 | < 0.0001 | 24/28 | 0.22 | 0.629 |

*In one participant, one 150-s block of the Regular–NoTask condition was replaced by the other block of the same condition because of artifacts. wMNE dWPLI, 38–42 Hz.*

### Reference

Colclough, G.L., Brookes, M.J., Smith, S.M., Woolrich, M.W., 2015. A symmetric multivariate leakage correction for MEG connectomes. NeuroImage 117, 439–448. https://doi.org/10.1016/j.neuroimage.2015.03.071
